# Phylogenetic and protein prediction analysis reveals the taxonomically diverse distribution of virulence factors in the *Bacillus cereus* group

**DOI:** 10.1101/2022.01.11.475806

**Authors:** Ming Zhang, Jun Liu, Zhenzhen Yin, Li Zhang

**Author notes:** Corresponding author, (ZY); (LZ).

## Abstract

*Bacillus cereus* is a food contaminant with widely varying enterotoxic potential of its virulence proteins. In this article, phylogenetic analysis of the whole-genome amino acid sequences of 41 strains, evolutionary distance calculation of the amino acid sequences of the virulence genes, and functional and structural prediction of the virulence proteins were performed to reveal the taxonomically diverse distribution of virulence factors. The genome evolution of the strains showed a clustering trend based on the coding virulence genes. The strains of *B. cereus* have evolved into non-toxic risk and toxic risk clusters with medium-high- and medium-low-risk clusters. The distances of evolutionary transfer relative to housekeeping genes of incomplete virulence genes were greater than those of complete virulence genes, and the distance values of HblACD were higher than those of nheABC and CytK among the complete virulence genes. Cytoplasmic localization was impossible for all the virulence proteins, and NheB, NheC, Hbl-B, and Hbl-L_1_ were extracellular according to predictive analysis. Nhe and Hbl proteins except CytK had similar spatial structures. The predicted structures of Nhe and Hbl mainly showed ‘head’ and ‘tail’ domains. The ‘head’ of NheA and Hbl-B, including two α-helices separated by β-tongue strands, might play a special role in Nhe trimers and Hbl trimers, respectively. The ‘cap’ of CytK, which includes two ‘latches’ with many β-sheets, formed a β-barrel structure with pores, and a ‘rim’ balanced the structure. The evolution of *B. cereus* strains showed a clustering tendency based on the coding virulence genes, and the complete virulence-gene operon combination had higher relative genetic stability. The beta-tongue or latch associated with β-sheet folding might play an important role in the binding of virulence structures and pore-forming toxins in *B. cereus*.

## Introduction

*Bacillus cereus* (*B. cereus*) is a gram-positive bacterium occurring ubiquitously in nature with widely varying pathogenic potential. *B. cereus*, the spores of which can survive at high temperatures and germinated vegetative cells of which can multiply and produce toxins under favorable conditions, is recognized as the most frequent cause of food-borne disease [1]. Its toxins cause two distinct forms of food poisoning, the emetic type (uncommon) and the diarrheal type (common). Diarrheal strains produce three enterotoxins, which belong to the family of pore-forming toxins: nonhemolytic enterotoxin (Nhe), hemolysin BL (Hbl), and cytotoxin K (CytK). Nhe comprises three proteins, NheA, NheB, and NheC, encoded by one operon containing one of three genes, namely, *nheA, nheB*, and *nheC*, respectively. Hbl consists of a single B-component (encoded by *hblA*) and two L-components, L_1_ (*hblC*) and L_2_ (*hblD*), all of which are essential for activity, with no individual or pairwise activity [2]. CytK (*cytK*) is a single-component toxin [3]. The genes that encode NheABC can be detected in nearly all enteropathogenic *B. cereus* strains, *hblBCD* can be detected in approximately 45% to 65% of such strains, and *cytK* is less prevalent [4,5]. *B. cereus* species which were compared on the basis of 16S rRNA (identity values >98%), were closely homologous to each other [6]. Some papers have reported that the species affiliation of *B. cereus* group strains, which could lead to an exchange of virulence plasmids between species, often does not match patterns of phylogenetic relatedness [7,8]. While the enterotoxins of *B. cereus* are chromosome-coded, the unique characteristics are observed for plasmids and are thus present throughout the *B. cereus* group [9]. Lapidus et al. (2008) reported a large plasmid with an operon encoding all three Nhe components in a *B. cereus* strain [10]. There is evidence that extensive gene exchange occurs between plasmids and the chromosome during the evolution of the *B. cereus* group [11]. Therefore, some genes encoded on plasmids can spread via horizontal gene transfer among *B. cereus* and the transfer of a single plasmid from one species to another [12]. Didelot et al. (2009) detected three phylogenetic groups (clades) in a study on the evolution of pathogenicity in the *B. cereus* group [13]. Later, seven major phylogenetic groups with ecological differences were identified in the *B. cereus* group [14]. A recent study suggested that nine phylogenetic clades of isolates may be better for assessing the risk of diarrheal foodborne disease caused by *B. cereus* group isolates [15]. These studies of virulence factors of *B. cereus* concern the evolutionary classification of virulence genes, and there have been few comparative analyses of the relative evolutionary distance of virulence genes and the prediction of virulence protein function and structure. In this study, the genome, virulence gene sequences and predicted virulence proteins of 41 *B. cereus* strains were comparatively analyzed. This work aims to examine the species diversity of the *B. cereus* group and the phylogenetic relationships among virulence factors, systematically evaluate the distribution of virulence genes, and comparatively analyze the structures and functions of virulence proteins.

## Materials and methods

### Characterization of *B. cereus* strains

Forty-one strains of *B. cereus* with complete chromosome-level assemblies in the National Center for Biotechnology Information (NCBI) database were selected for comparative analysis. We chose thirty-one pathogenic and ten nonpathogenic strains isolated from food, patients, the environment, and unknown sources and a control strain, *Sporolactobacillus terrae* (*T. Sporolactobacillus*), belonging to a different genus (details in Table 1). The sequences and annotation information of the stains were downloaded from the NCBI.

**Table 1.** The forty-one *B. cereus* strains and one control strain used in this study.

| Strain | Length<br>(Mb) | G+C<br>(%) | Completeness<br>(%) | Contamination<br>(%) | Origin | Enterotoxigenic<br>risk potential | Analysis of enterotoxigenic genes |  |  |
| --- | --- | --- | --- | --- | --- | --- | --- | --- | --- |
|  |  |  |  |  |  |  | *nheABC* | *hblACD* | *cytK* |
| B4264 | 5.42 | 35.30 | 98.99 | 0.00 | Pneumonia | H | + | + | + |
| ATCC14579 | 5.43 | 35.31 | 98.99 | 1.35 | Soil | H | + | + | + |
| FORC_013 | 5.68 | 35.20 | 98.99 | 0.00 | Foodborne | H | + | + | + |
| 03BB102 | 5.45 | 35.29 | 98.99 | 0.00 | Pneumonia | L | + | - | - |
| ATCC10987 | 5.43 | 35.52 | 98.99 | 0.00 | Soil | M | + | - | + |
| G9842 | 5.74 | 35.05 | 98.99 | 2.02 | Stool | M | + | + | - |
| AH820 | 5.59 | 35.31 | 98.48 | 0.00 | Periodontitis | H | + | + | + |
| F837/76 | 5.29 | 35.40 | 98.99 | 0.00 | Enteritis | M | + | + | - |
| Q1 | 5.51 | 35.50 | 98.99 | 0.34 | Nd | - | *AB* | - | - |
| CI | 5.49 | 35.27 | 98.99 | 1.01 | Anthrax | L | + | - | - |
| E33L | 5.84 | 35.17 | 98.99 | 0.00 | Carcass | M | + | - | + |
| AH187 | 5.60 | 35.52 | 98.99 | 0.00 | Nd | - | *AB* | - | - |
| NC7401 | 5.55 | 35.54 | 98.99 | 0.00 | Soil | - | *AB* | - | - |
| SGAir0263 | 6.47 | 34.91 | 98.99 | 0.34 | Air | M | + | + | - |
| SGAir0260 | 6.30 | 34.97 | 98.99 | 0.34 | Air | M | + | + | - |
| BHU1 | 5.20 | 35.10 | 98.99 | 0.00 | Soil | L | - | + | - |
| Co1-1 | 6.38 | 35.10 | 98.99 | 0.00 | Wastewater | L | - | + | - |
| ATCC4342 | 5.23 | 35.20 | 98.99 | 0.34 | Food | M | + | + | - |
| BDRD-Cer4 | 5.40 | 35.10 | 97.98 | 0.34 | Food | H | + | + | + |
| m1550 | 5.25 | 35.10 | 98.99 | 0.00 | Food | H | + | + | + |
| 95/8201 | 5.58 | 35.10 | 98.99 | 1.01 | Endocarditis | M | + | - | + |
| 172560W | 5.70 | 34.80 | 98.99 | 0.00 | Human | M | + | CD | + |
| Rock3-29 | 5.88 | 34.90 | 98.99 | 0.00 | Soil | - | AB | CD | - |
| Rock1-3 | 5.86 | 34.90 | 98.99 | 0.00 | Soil | - | AB | CD | - |
| Rock4-18 | 5.92 | 35.00 | 98.99 | 0.00 | Soil | - | AB | CD | + |
| Rock1-15 | 5.77 | 34.90 | 98.99 | 1.01 | Soil | H | + | + | + |
| BDRD- | 5.44 | 35.10 | 98.99 | 0.00 | Nd | H | + | + | + |
| ATCC10876 | 5.94 | 34.80 | 98.99 | 0.00 | Nd | H | + | + | + |
| Rock4-2 | 5.77 | 34.90 | 89.81 | 0.34 | Soil | H | + | + | + |
| BGSC6E1 | 5.73 | 35.00 | 98.99 | 0.00 | Nd | M | + | + | - |
| F65185 | 6.13 | 34.70 | 97.98 | 0.34 | Wound | M | + | AD | + |
| Rock3-28 | 6.04 | 35.75 | 96.46 | 0.34 | Soil | - | AB | CD | - |
| Rock3-42 | 5.20 | 35.20 | 98.99 | 0.00 | Soil | M | + | - | + |
| m1293 | 5.27 | 35.35 | 98.99 | 0.34 | Food | - | AB | - | - |
| BDRD- | 5.58 | 35.20 | 98.99 | 0.34 | Nd | - | AB | CD | - |
| AH676 | 5.59 | 35.00 | 98.99 | 1.01 | Soil | M | + | AD | + |
| AH1271 | 5.66 | 35.30 | 98.99 | 0.00 | Lamp | M | + | + | - |
| AH1272 | 5.79 | 35.20 | 97.47 | 1.35 | Amniotic | L | + | D | - |
| AH1273 | 5.79 | 35.25 | 96.93 | 1.35 | Blood | L | + | D | - |
| R309803 | 5.59 | 35.30 | 98.99 | 0.00 | Clinic | L | + | - | - |
| BDRD- | 5.57 | 35.25 | 98.99 | 0.00 | Nd | - | AB | - | - |
| Control-70- | 3.31 | 45.30 | 98.99 | 0.00 | Nd | - | - | - | - |

Nd: not determined; Enterotoxic: relative toxicity determined by gene expression; + or -: all or none, respectively; H, M, or L: three, two, or one type(s) of toxic genes, respectively. Enterotoxic risk potential: toxicity via the expression of the *nheABC*, *hblACD*, and *cytK* genes.

### Quality assessment of genomic sequences

The contamination and completeness of the metagenomic sequences were evaluated by CheckM software version v1.1.3 [16].

### Phylogenetic and average nucleotide identity (ANI) analysis

The first phylogenetic tree, based on whole-genome amino acid sequences of each strain, was constructed by using the CVTree4 webserver (http://cvtree.online/v4/prok/index.html), which constructs whole-genome-based phylogenetic trees without sequence alignment by using a composition vector (CV) approach, and the K-tuple length was 6 [17]. Every genome sequence was represented by a composition vector, which was calculated as the difference between the frequencies of k-strings and the prediction frequencies by the Markov model [18]. The shape and text content of the phylogenetic tree were modified by Molecular Evolutionary Genetics Analysis (MEGA-X version 10.2.2) [19]. In this study, the same genome sequence data were subjected to ANI analysis to verify the significance of the first phylogenetic tree. ANI analysis was performed using JSpeciesWS Online Service (http://jspecies.ribohost.com/jspeciesws/) as described by Richter et al. (2015) [20]. The distance matrix, which was calculated by the distance value (DV) using the formula DV=1-[ANIb value], was used to construct the second phylogenetic tree, which was generated from the resulting Newick format file using Njplot [21]. The formula was balanced using the mean value method and was subjected to calculation using DrawGram in the PHYLIP package version 3.695 [22].

### Multilocus sequence analysis (MLSA)

A total of forty-one strains containing gene sequences, which were downloaded from the NCBI, were found and further analyzed for the presence of seven housekeeping and three enterotoxin genes. The housekeeping genes adenylate kinase (*adk*), catabolite control protein A (*ccpA*), glycerol uptake facilitator protein (*glpF*), glycerol-3-phosphate transporter (*glpT*), pantoate-beta-alanine ligase (*panC*), phosphate acetyltransferase (*pta*), and pyruvate carboxylase (*pyc*) were chosen to calculate the basic evolutionary distances of the species. These housekeeping genes, scattered across the entire chromosome, are suitable for MLSA [23]. The types of enterotoxin genes (*nhe, hbl*, and *cytK*) were divided into different groups, and the base sequences were concatenated for further MLSA. Thus, rearrangement of genes was unnecessary because the order of the genes within the operons was conserved in all strains. The distances of concatenated genes were calculated in MEGAx using the maximum likelihood (ML) algorithms, which are based on the Tamura-Nei model with a discrete gamma distribution [19]. The model applied for MLSA of DVs was the ideal substitution model according to the ‘find best DNA/Protein models’ function [24]. The housekeeping genes were of the same length in all strains, as were the different virulence genes. The same settings for the calculation of all phylogenetic DVs were used to ensure comparability of the results. We calculated the relative changes in genetic DVs between the virulence genes, which were concatenated housekeeping genes minus the simple housekeeping gene, representing the change in virulence gene transfer.

### Prediction of virulence protein function and structure

SMART software (http://smart.embl-heidelberg.de/), which is a simple modular architecture research tool, was used to predict the domain architecture of the virulence proteins in this study [25]. PSORT (http://www.psort.org/psortb2) and TMHMM (http://www.cbs.dtu.dk/services/TMHMM) software were employed to predict the subcellular location and transmembrane helices of virulence proteins, respectively [26,27]. The SIGNALP-5.0 (http://www.cbs.dtu.dk/services/SignalP/) and SWISS-MODEL (http://swissmodel.expasy.org) servers and AlphaFold v2.1.1 (https://github.com/deepmind/alphafold) were used to predict the signal peptide cleavage and three-dimensional (3-D) structures of the enterotoxin proteins, respectively [28–31]. The amino acid sequences of the virulence proteins analyzed were submitted in FASTA format. To predict structure, we performed homology modeling to generate 3-D virulence protein structures.

## Results

### General genome characteristics and quality assessment of sequences

A summary of the features of the forty-one genomes of *B. cereus* and the control genome of the closely related species *T. sporolactobacillus* is provided in Table 1. The genome sizes of *B. cereus* strains varied from 5.20 to 6.47 MB. The G+C contents of the forty-one genomes ranged from 34.70% to 35.75%. Compared with the control genome from *T. sporolactobacillus*, the genomes of *B. cereus* were much larger and had lower G+C contents. The contamination and completeness of the sequences were 0-2.02% and 89.81%-98.99%, respectively (shown in Table 1). The results suggested that these sequences are of high quality and have low contamination (values <2.02%) and high completeness (values >89.81%); thus, they were appropriate for analysis. In this study, the strains originated from food (5/41), the clinic (12/41), the environment (17/41), and undetermined sources (7/41). The enterotoxic risk potential based on the virulence genes of forty-one *B. cereus* strains is listed in Table 1. Enterotoxicity, which was reflected by virulence gene numbers, was categorized into levels of three types (10/41), two types (14/41), one type (7/41), and no types (10/41) levels. The genes detected as enterotoxic were *nheABC* (29/41), *hblACD* (19/41), *cytk* (18/41), *nheAB* (10/41), *hblCD* (6/41), *hblAD* (2/41), and *hblD* (2/41).

### Phylogenetic analysis based on whole amino acid sequences

Two whole-genome-based methods were used to construct phylogenetic trees. The first phylogenetic tree was constructed with the CV method using the whole amino acid sequences of forty-one *B. cereus* group strains and the outgroup species *T. sporolactobacillus* [32]. To ensure the accuracy of the results, we added the inbuilt sequence AH1273 and the sequence of *T. sporolactobacillus* (control) from the webserver database (Fig 1). According to enterotoxic risk potential, the forty-one strains of *B. cereus* had evolved into five distinct clusters, which were likely risk regions I, IV and V and nonrisk regions II and III. However, there were individual nonconformities, such as nontoxicity of BDRD ST196 in region V. Region I was dominated by medium- and high-risk strains (15/17) but also included two low-risk strains (BHU1 and CO1-1), region II and III included only nonrisk strains (9/9), and regions IV and V were dominated by medium- and low-risk strains (13/15) but also included AH820 (high-risk strain) and BDRD ST196 (nonrisk strain), respectively.

**Fig. 1.**
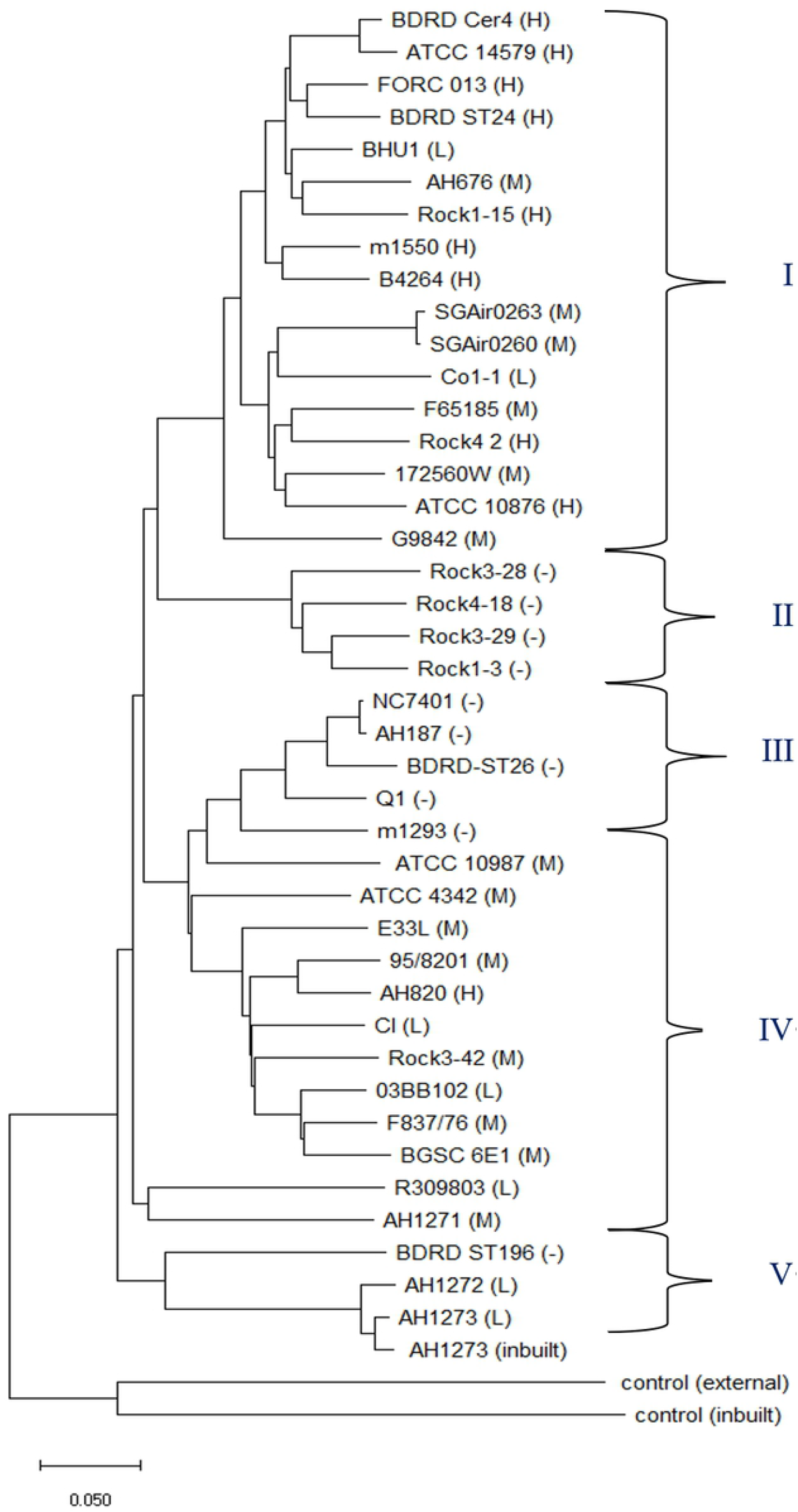
Phylogenetic relationships of the amino acid sequences of forty-one *B. cereus* strains and one external control strain used in this study. Two additional inbuilt strains in the software were used as internal controls. H, M, L, and “-” symbols indicate high (three types of virulence genes), middle (two types of virulence genes), low (one type of virulence gene), and no enterotoxic risk potential.

To verify the above results and obtain more accurate molecular evolutionary relationships, we established a second phylogenetic tree based on ANI analysis (Fig 2). According to enterotoxic risk potential, the forty-one strains of *B. cereus* had evolved into six distinct regions: likely risk regions A, C, E, and D_2_ and nonrisk regions B and D_1_. The two phylogenetic trees were similar in terms of the regions where enterotoxic risk was likely. Region A, which was dominated by medium-high-risk strains (15/17) but also included two low-risk strains (BHU1 and CO1-1), was the same as region I. Regions B and D_1_, which included only nonrisk strains (9/9), were the same as regions II and III. Regions C, D_2_ and E, which were also dominated by medium- and low-risk strains (13/15) but included AH820 (high-risk strain in C) and BDRD ST196 (nonrisk strain in E), were the same as regions IV and V.

**Fig. 2.**
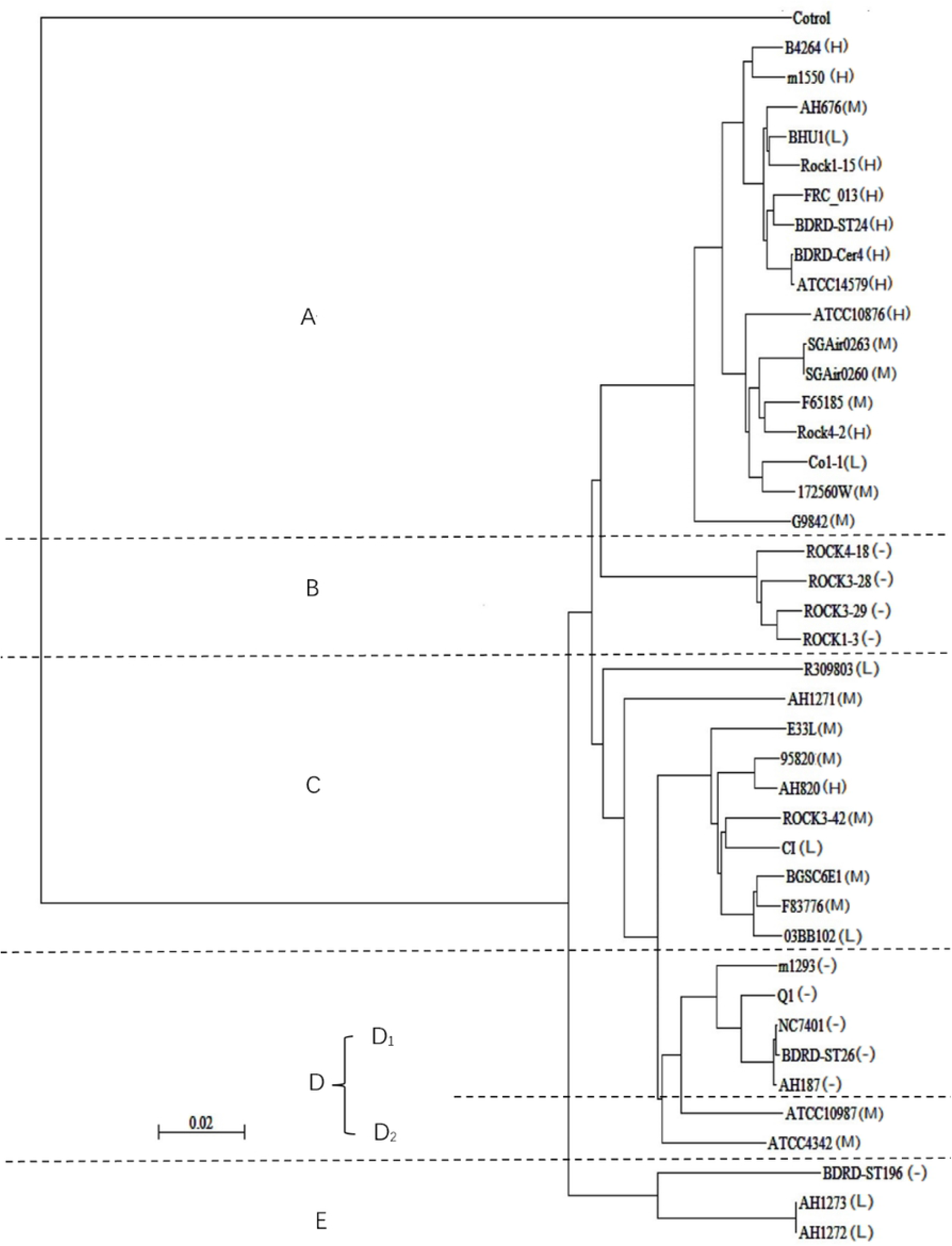
ANI analysis of the phylogenetic relationships of the forty-one *B. cereus* strains and one external control strain used in this study. H, M, L, and “-” have the same meanings as in Table 1.

### Phylogenetic distance analysis based on concatenated housekeeping and virulence genes

To analyze the evolution and phylogenetic relationships of virulence gene transfer in relation to DVs, the *nheABC, hblACD*, and *cytK* genes of the forty-one strains, which need to be compared to the housekeeping genes of the strains, were studied. To this end, we concatenated the sequences of virulence proteins from the strains and seven housekeeping proteins (Adk-CcpA-GlpF-GlpT-PanC-Pta-Pyc) from the *B. cereus* core genome. The genetic DV of virulence gene transfer was evaluated by calculating the average difference in the phylogenetic DV of the ATCC14579 strain compared with forty other strains. The hblD and hblAD virulence proteins, which were observed in only two strains, were excluded. As shown in Table 1, the relative genetic DVs were calculated for nheAB (10/41), hblCD (6/41), nheABC (29/41), hblACD (19/41), and CytK (18/41). As shown in Fig 3, the average evolutionary DVs of virulence gene transfer from high to low were 0.015 (*nheAB*), 0.012 (*hblCD*), 0.005 (*hblACD*), 0.003 (*nheABC*) and 0.001 (*cytK*). The DVs of incomplete virulence genes (*nheAB* and *hblCD*) were higher than those of complete virulence genes (*nheABC, hblACD*, and *CytK*). The average evolutionary DV of *hblACD* was the highest and that of *cytK* was the lowest among the complete virulence genes.

**Fig. 3.**
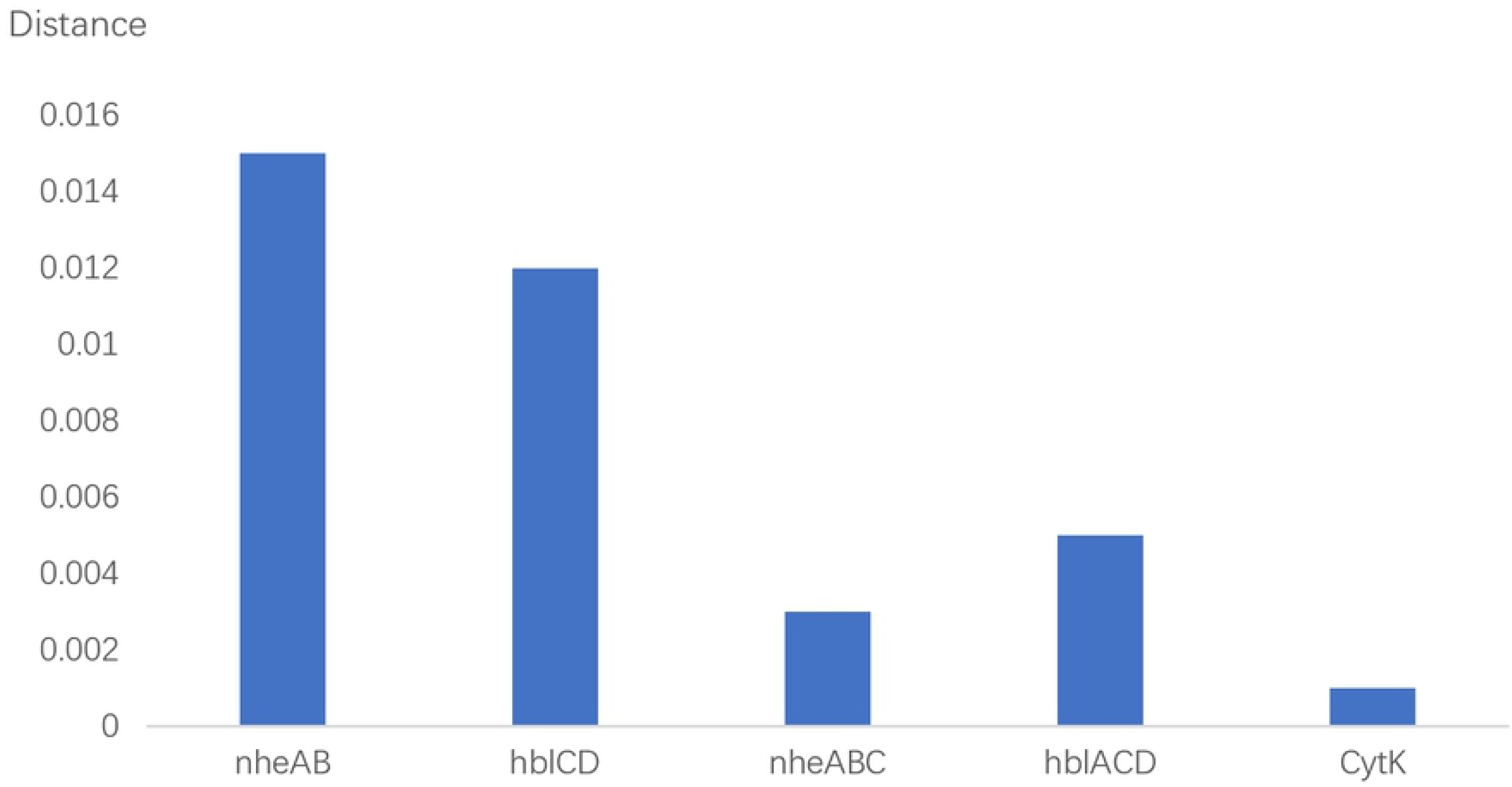
The average difference in the phylogenetic distance values of the ATCC14579 strain compared with forty other strains for virulence genes plus housekeeping genes and housekeeping genes examined by MLSA.

### Comparative prediction analysis of the function and structure of virulence proteins

As shown in Table 2, we obtained the scores of the seven virulence proteins for subcellular localization prediction. The scores of NheB, NheC, Hbl-B, and Hbl-L_1_ were all 9.73, and that of CytK was 9.98, all consistent with extracellular localization. The localization of NheA and Hbl-L_2_ was unknown because the scores were all lower in the cytoplasmic membrane (3.33/4.6), cell wall (3.33/2.48), and extracellular space (3.33/2.92), making it impossible for the virulence proteins to appear in the cytoplasm. As shown in Fig 4, the amino acid sequences of Nhe, Hbl and CytK contained N-terminal signal peptides for secretion (sequences < 31), NheB and Hbl-L_1_ had two helices, which were transmembrane region sequences 235-257/267-286 and 239-261/268-290, and NheC had only one helix, of which the transmembrane region was 228-250, but the others had none. The virulence protein cleavage sites of all strains were in the sequence 30-32 with 0.93-0.99 likelihood levels, except NheA, for which the site was in the 26-27 sequence (0.81 likelihood level). The signal peptide start-end was between 1 and 31 sequences but not found for Hbl-B and Hbl-L_1_ were not found, and the domain start-end was between 35 and 329 sequences.

**Fig. 4.**
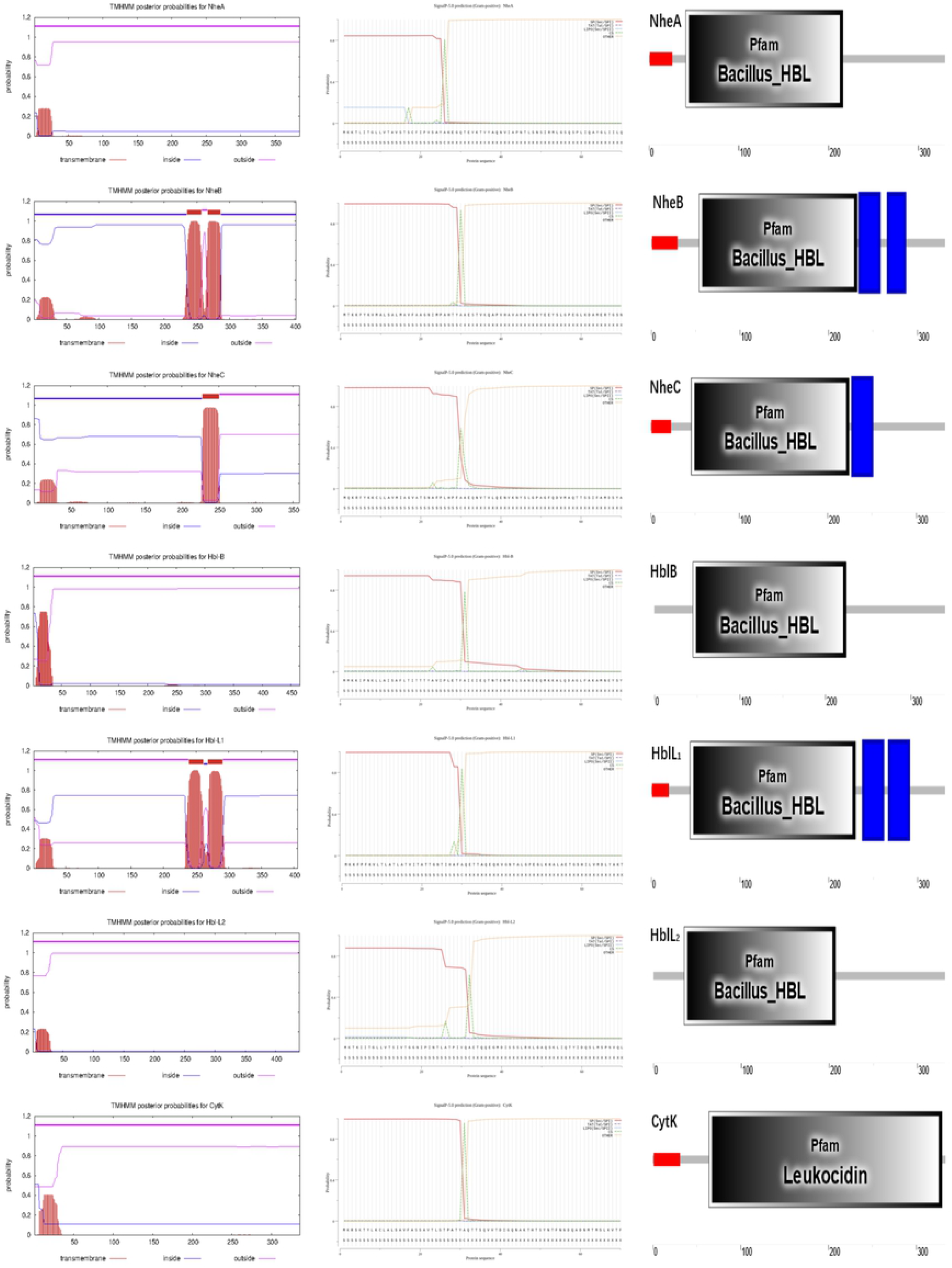
The locations of transmembrane helices, cleavage sites, signal peptides, and domain start-ends were predicted by TMHMM, SignaIP, and SMART software with the ATCC14579 strain. The symbols are 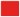signal peptide, 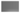domain, and 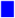transmembrane region.

**Table 2.** Subcellular localization, transmembrane helix and region, signal peptide, and domain prediction results of the virulence proteins of *B. cereus* strains.

| Name | Final localization | Final score | Scores for different localizations |  |  |  | Helix number | Transmembrane region | Cleavage site | Signal peptide likelihood | Signal peptide start-end | Domain start-end |
| --- | --- | --- | --- | --- | --- | --- | --- | --- | --- | --- | --- | --- |
|  |  |  | Cytoplasmic membrane | Cell wall | Extracellular | Cytoplasmic |  |  |  |  |  |  |
| NheA | Unknown | 3.33/4.60 | 3.33/4.60 | 3.33/2.48 | 3.33/2.92 | 0.00 | 0 | - | 26 and 27 | 0.81±0.07 | 1-26 | 41-216 |
| NheB | Extracellular | 9.73 | 0.09 | 0.18 | 9.73 | 0.00 | 2 | 235-257;267-286 | 30 and 31 | 0.99±0.01 | 1-30 | 54-232 |
| NheC | Extracellular | 9.73 | 0.09 | 0.18 | 9.73 | 0.00 | 1 | 228-250 | 30 and 31 | 0.98±0.02 | 1-23 | 46-224 |
| Hbl-B | Extracellular | 9.73 | 0.09 | 0.18 | 9.73/9.72 | 0.00 | 0 | - | 31 and 32 | 0.96±0.04 | - | 45-224 |
| Hbl-L<sub>1</sub> | Extracellular | 9.73 | 0.09 | 0.18 | 9.73 | 0.00 | 2 | 239-261;268-290 | 30 and 31 | 0.98±0.01 | 1-20 | 44-230 |
| Hbl-L<sub>2</sub> | Unknown | 3.33/4.60 | 3.33/4.60 | 3.33/2.48 | 3.33/2.92 | 0.00 | 0 | - | 32 and 33 | 0.93±0.05 | - | 35-207 |
| CytK | Extracellular | 9.98 | 0.00 | 0.02 | 9.98 | 0.00 | 0 | - | 31 and 32 | 0.99±0.01 | 1-31 | 64-329 |

Examination of the phylogenetic tree constructed using the Hbl, Nhe, and CytK sequences (Fig 5) showed that NheA and Hbl-L_2_ as well as NheBC and Hbl-L_1_ were more closely related to one another than to the other components, and CytK was the least evolutionarily related. This result was also reflected in the evaluation parameters of the 3-D enterotoxin protein structures. As shown in Table 3, the closest template of NheB and NheC was Hbl-L_1_ (sequence identity of 40.82% and 36.83%, respectively), and that of Hbl-L_2_ was NheA (24.95%). That of CytK was alpha-hemolysis (30.39%), as expected, with considerable amino acid sequence homology to *S. aureus* leukocidin [33]. The templates of NheA, Hbl-B, and Hbl-L_1_ were included in the SWISS-MODEL server with high sequence identity (97.22%, 71.99%, and 99.73%, respectively). The sequence coverage and range of all structures were 0.71-0.93 and 33-439, respectively, with GMQE evaluation values above 0.5, which indicated reliable model construction. Each residue is allotted a reliability score between 0 and 1, indicating the expected resemblance to the native structure. Higher numbers represent higher reliability of the residues [34]. To verify the above results (the sequence identity of Hbl-L_2_ was 24.95%) and obtain the predicted Nhe-trimer and Hbl-trimer structures, we used AlphaFold software for secondary structural prediction. The results were acceptable, the predicted local-distance difference test (plDDt) values of monomers were 81.90-94.14, and the scores of Nhe trimers and Hbl trimers were 0.68 and 0.36 (ipTM+pTM), respectively [30,31].

**Fig. 5.**
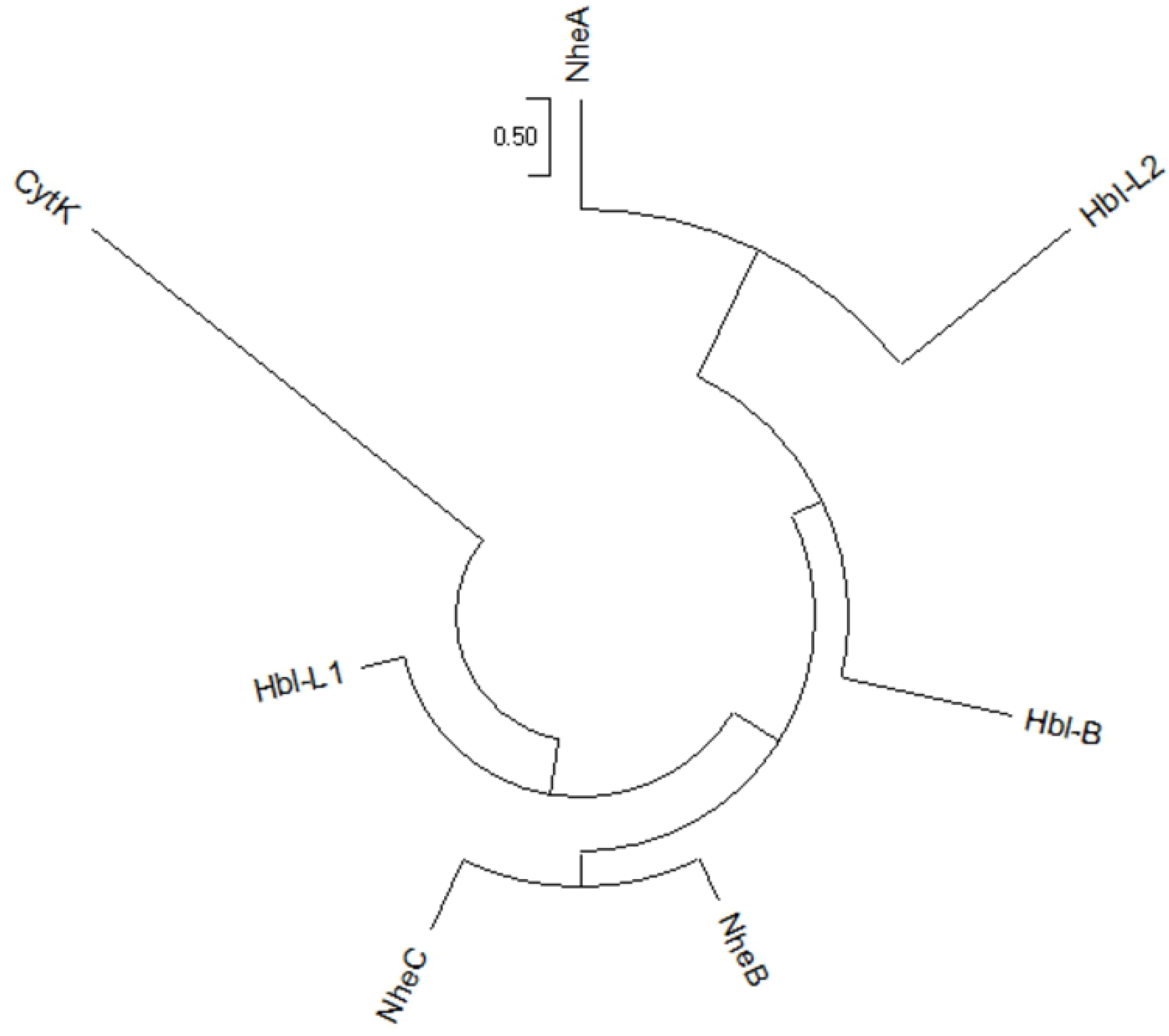
The phylogenetic tree of Hbl-B, Hbl-L_1_, Hbl-L_2_, NheA, NheB, NheC, and CytK component sequences with the ATCC14579 strain.

**Table 3.**
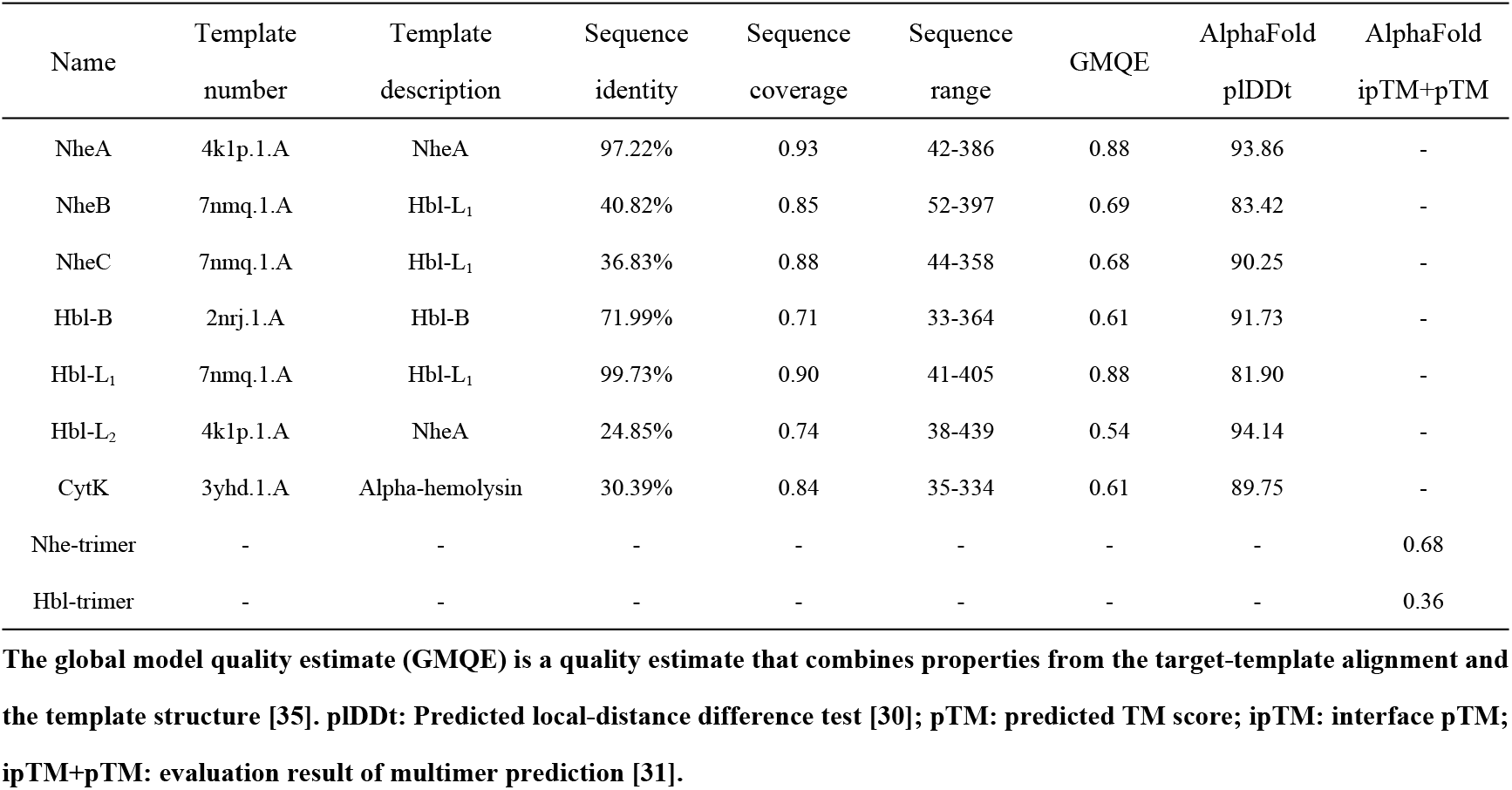
The evaluation parameters of the 3-D enterotoxin protein structures predicted by the SWISS-MODEL server and AlphaFold software with the ATCC14589 strain.

| Name | Template number | Template description | Sequence identity | Sequence coverage | Sequence range | GMQE | AlphaFold pLDDt | AlphaFold ipTM+pTM |
| --- | --- | --- | --- | --- | --- | --- | --- | --- |
| NheA | 4k1p.1.A | NheA | 97.22% | 0.93 | 42-386 | 0.88 | 93.86 | - |
| NheB | 7nmq.1.A | Hbl-L<sub>1</sub> | 40.82% | 0.85 | 52-397 | 0.69 | 83.42 | - |
| NheC | 7nmq.1.A | Hbl-L<sub>1</sub> | 36.83% | 0.88 | 44-358 | 0.68 | 90.25 | - |
| Hbl-B | 2nrj.1.A | Hbl-B | 71.99% | 0.71 | 33-364 | 0.61 | 91.73 | - |
| Hbl-L<sub>1</sub> | 7nmq.1.A | Hbl-L<sub>1</sub> | 99.73% | 0.90 | 41-405 | 0.88 | 81.90 | - |
| Hbl-L<sub>2</sub> | 4k1p.1.A | NheA | 24.85% | 0.74 | 38-439 | 0.54 | 94.14 | - |
| CytK | 3yhd.1.A | Alpha-hemolysin | 30.39% | 0.84 | 35-334 | 0.61 | 89.75 | - |
| Nhe-trimer | - | - | - | - | - | - | - | 0.68 |
| Hbl-trimer | - | - | - | - | - | - | - | 0.36 |

The global model quality estimate (GMQE) is a quality estimate that combines properties from the target-template alignment and the template structure [35]. pLDDt: Predicted local-distance difference test [30]; pTM: predicted TM score; ipTM: interface pTM; ipTM+pTM: evaluation result of multimer prediction [31].

Due to the sequence similarity of NheB and NheC with Hbl-B, homology models based on the Hbl-B structure were established. As shown in Fig 6, NheA and Hbl-B had highly similar structures (Fig 6a), and the NheA, NheB, NheC, HBl-B, HBl-L_1_, and HBl-L_2_ structures showed that there were two main domains, a ‘head’ and ‘tail’ (Figs 6b-d, 6f-h, and 7A-F). The main body of the structure was formed by the ‘tail’ domain, which consisted of five major helices, and the ‘head’ domain of NheA included two long α-helices separated by β-tongue strands (Figs 6b and 7A). Multiple β-tongue strands were detected in Hbl-B (Fig 7D) but are not shown in Fig 6f because of the prediction method. Another difference was the ‘head’ of Hbl-L_2_, possibly related to the low sequence identity (Figs 6h and 7F). The ‘latch’ with many β-sheets of CytK folded the ‘cap’ domain, which was the toxic area (Figs 6e and 7G). The amino ‘latch’, which included a short helix in all known pore structures, was observed on the top of the conformation, which extended into the pore to form a β-barrel and was folded into a stranded antiparallel β-sheet in the monomer. Although the amino ‘latch’ protrudes and interacts with the adjacent protomer in the pore, it is located at the edge of the β-sheet of the ‘cap’ region [36]. The ‘rim’ domain, which was composed of three strands of short β-sheets, formed the main body of the balanced structure. The trimers of Nhe and Hbl were horizontally arranged. The Hbl trimer (arranged in the sequence B, -L_1_, -L_2_) was more similar than the Nhe trimer (arranged in the sequence A, B, C) based on the structural features. The β-tongue strands of Hbl-B and NheA might play an important structural and functional role in the formation of trimers.

**Fig. 6.**
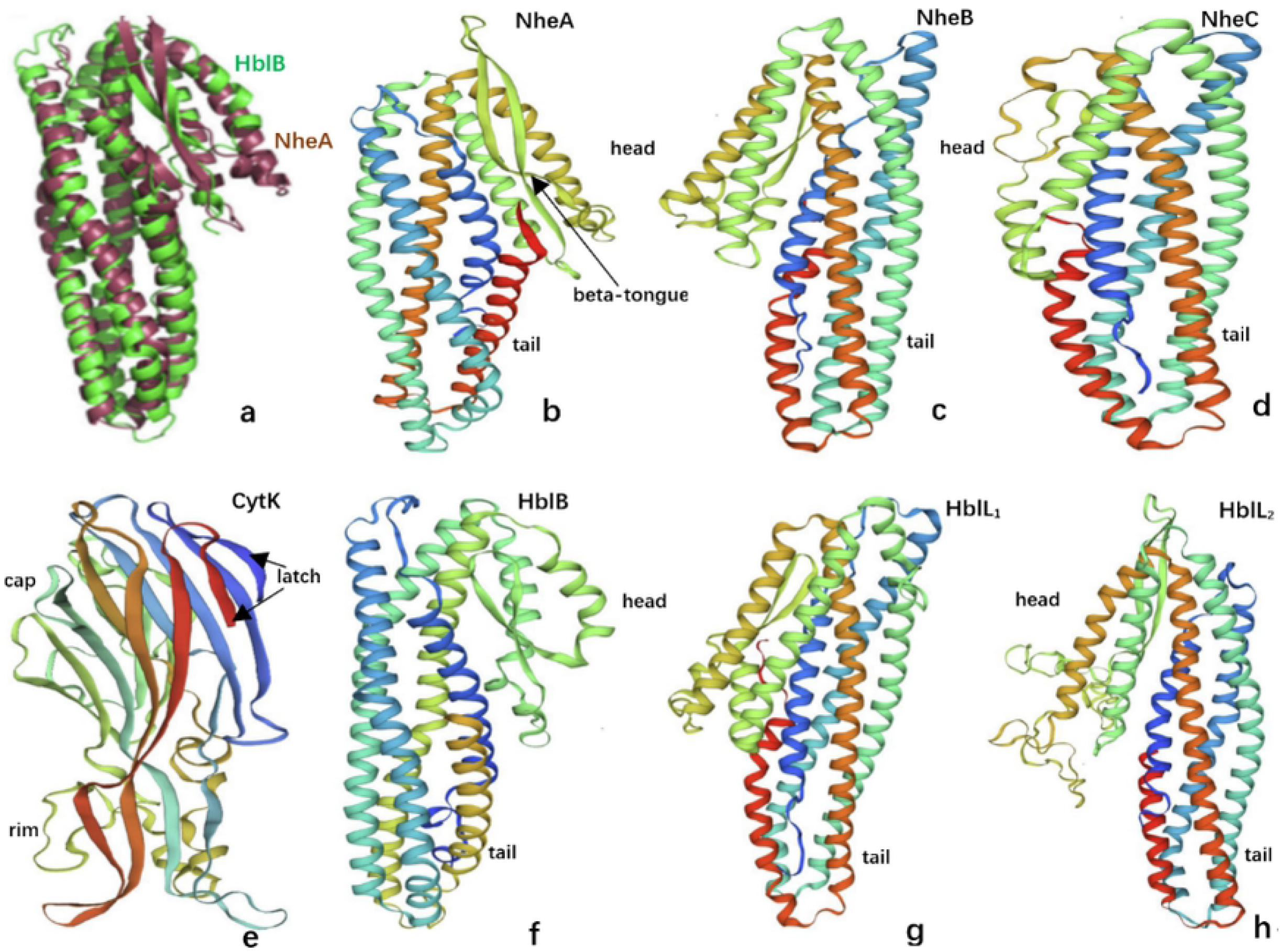
Overview of the structure predicted by the SWISS-MODEL server. a shows the superposition of the structures of Hbl-B (green) and NheA (burgundy) [37]. b, c, d, f, g, and h show the structures of NheA, NheB, NheC, Hbl-B, Hbl-L_1_ and Hbl-L_2_, which are annotated with the ‘head’ and ‘tail’, respectively. b shows a beta-tongue in the ‘head’ region. e shows the structure of CytK, which is annotated with ‘latch’, ‘cap’ and ‘rim’.

**Fig. 7.**
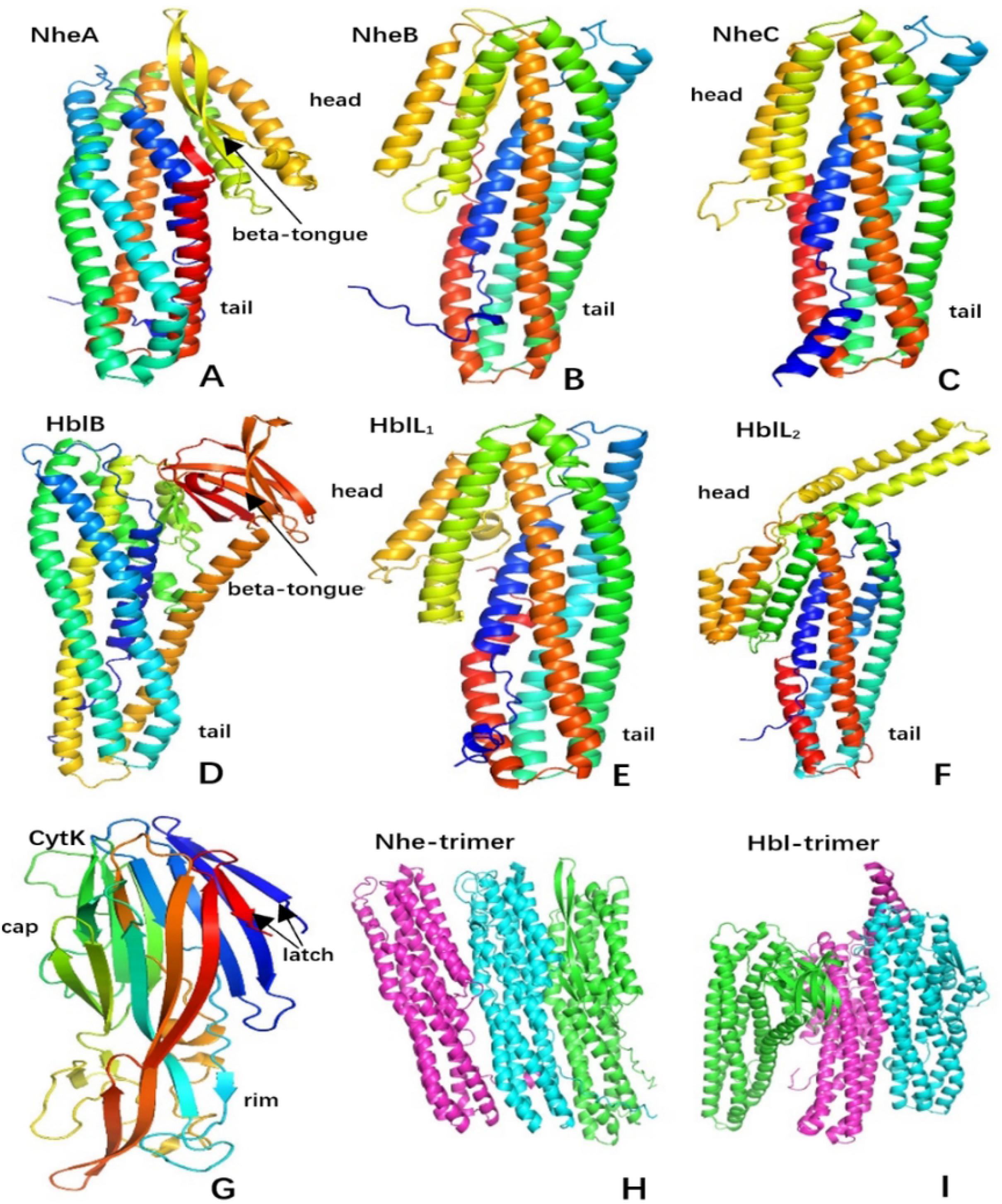
Overview of the structure predicted by AlphaFold software. A, B, C, D, E, and F are the structures of NheA, NheB, NheC, Hbl-B, Hbl-L_1_, and Hbl-L_2_, which are annotated with the ‘head’ and ‘tail’, respectively. A and D show beta-tongues in the ‘head’ region. G is the structure of CytK, which is annotated with ‘latch’, ‘cap’ and ‘rim’. H and I show the trimers of the structures of NheA and Hbl-B (green), NheB and Hbl-L_2_ (wathet), and NheC and Hbl-L_1_ (pink).

## Discussion

In this study, forty-one strains of the *B. cereus* group were subjected to phylogenetic analyses based on whole amino acid sequences. Enterotoxicity, which was evaluated on the basis of *nheABC, hblACD*, and *cytK* gene expression, was classified into levels of three types, two types, one type, and no types. In terms of evolutionary relationships, clusters of virulence and nonvirulence gene strains were evident, and the regional distribution of the number of types of virulence genes was also presented, further confirmed by ANI-based phylogenetic analyses. We found that the two phylogenetic trees were similar. All non-toxic-risk strains were concentrated in two clusters, and all but two of the medium-high- and medium-low-toxic-risk strains formed clusters. The results suggest the possibility of virulence gene transfer, which may be related to frequent exchange of pathogenicity factors during *B. cereus* virulence evolution, including so-called probiotic or nonpathogenic species [12]. The inconsistent evolutionary distribution of individual virulence genes may be due to other factors, which needs further study.

The bacillus hemolytic and nonhemolytic enterotoxin family of proteins consists of several bacillus enterotoxins, which can cause food poisoning in humans [38]. Hemolytic BL and cytotoxin K (encoded by *hblACD* and *cytK*) and nonhemolytic enterotoxin (encoded by *nheABC*) represent the significant enterotoxins produced by *B. cereus*. Cardazzo et al. (2008) detected horizontal gene transfer in the evolution of enterotoxins within the *B. cereus* group [39]. Our MLSA results showed that in the process of toxin molecular evolution, two toxic-type genes had a more significant effect in relation to DVs than three toxic-type genes. The results suggested that the complete virulence-gene operon combination has higher relative genetic stability. The DV of hemolysin Bl was greater than that of nonhemolytic cytotoxin K. *nheABC*, which was responsible for most of the cytotoxic activity of *B. cereus* isolates, showed stable, strictly vertical inheritance [40]. In contrast to *hbl*, duplication or deletion of *nhe*, which was almost exclusively transmitted vertically, was rarely observed, and *cytK*, a one-type gene, had the highest relative genetic stability [12].

Currently, it is commonly accepted that the toxicity potential of *B. cereus* is not driven by enterotoxin gene types because the expression of enterotoxin genes is highly complex and probably strain-specifically affected by transcription, posttranscriptional and posttranslational modification [41–43]. Both Nhe and Hbl are three-component cytotoxins composed of binding components A and B and two lytic components B, C and -L_1_ -L_2_, with all three subunits acting synergically to cause illness. The amino acid sequences of all Nhe, Hbl and Cytk components containing N-terminal signal peptides indicated toxin secretion via the secretory translocation pathway. The final positions of Hbl-B, Hbl-L_1_, NheB, NheC, and CytK were all extracellular and did not appear in the cytoplasm, and the two transmembrane regions of NheB and Hbl-L_1_ might be responsible for transporting the assembled three-component cytotoxins across the membrane to complete the toxic effect. Dietrich et al. (2021) found that the factor triggering enterotoxin production under simulated intestinal conditions by various cell lines from different organisms and compartments was independent of cell differentiation [44]. Similarities were found when predicted transmembrane helices were compared. NheA and Hbl-B had no such helices, and NheB and Hbl-L_1_ had two that may play an important role in molecular docking and transmembrane activities. Furthermore, the Nhe components seem to be additionally processed in the extracellular space after separation from the signal peptide for secretion [44]. The difference is that NheC had one such component and Hbl B had none, which may strengthen the secretion of the Nhe protein.

The Nhe and Hbl proteins share sequence similarities, both between the three components of each complex and between the two enterotoxin complexes [9]. The structural and functional properties were consistent with those of the superfamily of pore-forming cytotoxins of Hbl and Nhe [45–47]. The NheA, NheB, NheC, HBl-B, HBl-L_1_, and HBl-L_2_ structures showed two main domains, a ‘head’ and ‘tail’. The ‘heads’ of NheA and Hbl-B, including two α-helices separated by β-tongue strands, play a special role in Nhe trimers and Hbl trimers, respectively. Upon contact with lipids, cell membranes or detergents, the protein oligomerizes and forms ring-shaped structures acting as transmembrane pores [48,49], and the hydrophobic β-tongue is assumed to be inserted into the membrane first [50]. It is worth noting that NheB, NheC, Hbl-L_1_, and Hbl-L_2_ had few or no β-strands in the ‘head’, which were either responsible for conformational changes of NheA and Hbl-B or for the stabilization of the ‘head’ domain [46] or might lead to reduced toxicity or only ligand function [37]. The difference was reflected in the triplet prediction results, i.e., a significant difference in structural arrangement compactness between Nhe trimers (noncompact type) and Hbl trimers (compact type). Ganash et al. (2013) speculated that the Nhe trimer requires interaction with unknown proteins of an additional function [37]. A specific binding order of the three Nhe and Hbl components is also necessary for pore formation [51,52]. We found that NheB and Hbl-L_1_ were close to NheA and Hbl-B in the predicted trimer structure. Didier et al. (2016) found that NheA is important for attaching to cell-bound NheB and NheC and that NheB is the main interaction partner of NheA [53], and a further correlation was found for the amounts of Hbl B and Hbl L_1_ [42]. Cytotoxin K is a single protein with β-barrel pore-forming toxin in contrast to the tripartite toxin complexes Hbl and Nhe. The CytK structure, which exhibits two ‘latches’ with many β-sheets folded beside the ‘cap’ domain forming a β-barrel, was the pore structure on top of the conformation. The ‘rim’ region, which was folded into a three-stranded antiparallel β-sheet, balanced the structure in the monomer. The predicted structure revealed that CytK was likely to belong to the leukocyte toxin family. These monomers diffuse to target cells and are attached to them by specific receivers [54], which are lipids and proteins that cause lysis of red blood cells by destroying their cell membrane [55].

## Conclusion

In this study, we describe the molecular evolution, function and structural diversity of virulence factors in the *B. cereus* group. The evolution of *B. cereus* strains showed a clustering trend based on the coding virulence genes. The complete virulence gene operon combination had higher relative genetic stability than the incomplete operon. The two α-helices in the ‘head’ of the NheA and Hbl-B structures, which are separated by β-tongue strands, and two ‘latches’ with many β-sheets folded beside the ‘cap’ of the CytK structure might play a special role in the binding of virulence structures and pore-forming toxins in *B. cereus*. Overall, the exact mechanism by which *B. cereus* causes diarrhea remains unknown, but our results provide helpful information for better understanding the taxonomically diverse distribution of virulence factors in the *B. cereus* group.

## Abbreviations

ANI: average nucleotide identity
MLSA: multilocus sequence analysis
Nhe: nonhemolytic enterotoxin
Hbl: hemolysin BL
CytK: cytotoxin K
DV: distance value
CV: composition vector
GMQE: global model quality estimate
NCBI: National Center for Biotechnology Information
*adk*: adenylate kinase
*ccpA*: catabolite control protein A
*glpF*: glycerol uptake facilitator protein
*glpT*: glycerol-3-phosphate transporter
*panC*: pantoate-beta-alanine ligase
*pta*: phosphate acetyltransferase
*pyc*: pyruvate carboxylase

## Author Contributions

Conceptualization: Ming Zhang

Formal analysis: Jun Liu

Funding acquisition: Ming Zhang

Methodology: Zhenzhen Yin

Resources: Ming Zhang

Writing-original draft: Ming Zhang

Writing-reviewing and editing: Li Zhang

